# Physiologically Based Pharmacokinetic Modeling of mRNA-Encoded Therapeutics: A Multiscale Framework for LNP and Antibody Trafficking in Mice

**DOI:** 10.64898/2025.12.20.695667

**Authors:** Elio Campanile, Elisa Pettinà, Stefano Giampiccolo, Lorena Leonardelli, Luca Marchetti

## Abstract

Antibody-based therapeutics have revolutionized disease treatment, and recent advances in messenger RNA (mRNA) technologies have opened new opportunities for their intracellular production. In particular, in vitro–transcribed mRNA encapsulated in lipid nanoparticles (LNPs) enables targeted delivery to specific cells, where it can enable the synthesis of therapeutic antibodies with prolonged half-lives in a cost-effective manner. Despite rapidly growing experimental data, a modeling framework that integrates mRNA delivery, intracellular expression kinetics, and whole-body antibody disposition remains unavailable. To address this gap, we extended a Physiologically Based Pharmacokinetic model with a novel multiscale layer describing mRNA trafficking, cellular uptake, translation, and degradation. The integrated model was calibrated and validated using five datasets of mRNA-based cancer therapeutics, demonstrating strong predictive performance for the biodistribution of mRNA-encoded antibodies. The newly introduced mRNA layer, while minimally parameterized, effectively represents complex intracellular and systemic processes, enabling quantitative investigation of antibody biodistribution, optimization of dose scheduling, and providing an initial framework for future exploration of how LNP–mRNA formulation influences delivery and pharmacokinetics.

## Introduction

Monoclonal antibodies (mAbs) and bispecific antibodies (bsAbs) have become central pillars of modern biologic therapeutics, with applications spanning autoimmune, infectious, and inflammatory diseases as well as targeted cancer therapy^1–5^. mAbs have demonstrated remarkable clinical efficacy across diverse disease areas due to their high antigen specificity and long systemic persistence^6^. bsAbs further expand these therapeutic capabilities by simultaneously engaging two distinct antigens, enabling mechanisms such as enhanced target selectivity or coordinated immune-cell recruitment^5^.

Despite their clinical potential, mAbs and bsAbs present significant pharmacokinetic (PK) and manufacturing challenges, such as short half-life, costly and complex production processes, and potential immunogenicity^7–9^. One approach to address some of these limitations is based on messenger RNA (mRNA)-based therapeutics. Following the unprecedented success of mRNA vaccines against COVID-19, mRNA technologies have rapidly advanced toward therapeutic applications, enabling the *in situ* production of complex proteins directly within the patient’s own cells^10,11^. mRNA constructs encoding antibodies, usually encapsulated in lipid nanoparticles (LNPs) to prevent degradation and favor delivery, aim to generate functional mAbs or bsAbs endogenously, potentially streamlining manufacturing through scalable nucleic-acid production while improving pharmacokinetic control and tissue distribution. Moreover, the gradual intracellular synthesis of the encoded antibody may mitigate immunogenicity relative to bolus administration of recombinant proteins. Among potential target tissues, liver hepatocytes are particularly efficient in mRNA uptake and protein expression, making them optimal cellular factories for the endogenous production and systemic secretion of antibodies directed against tumor-associated antigens^12,13^. In the present study, we therefore focus our analysis to conventional liver-tropic LNP formulations, for which functional delivery is expected to be predominantly hepatocyte-driven.

Despite their remarkable therapeutic potential, the clinical applicability of mRNA-LNP antibodies faces several challenges, mainly arising from the limited knowledge of key intracellular biological steps and how to tie them to macroscopic, often qualitative measurements^14–16^. For example, it is well documented that, in cells, the endosomal escape of mRNAs is a rate-limiting step^17^, but the impossibility of gaining quantitative mRNA data from liver tissue in vivo and having to rely exclusively on time series in plasma, makes it extremely difficult to devise equations that are informative to such small time scales. In parallel, the antibodies produced from these constructs—ranging from mAbs to bsAbs—can differ greatly in terms of pharmacokinetics, target engagement modalities, and clearance profiles, complicating the development and optimization process^4,5^.

This is where mathematical modeling becomes a powerful tool. Supported by the FDA’s latest guidelines on Model-Informed Drug Development (MIDD)^18^, modeling approaches are increasingly used to inform decision-making, especially for complex biologics and novel modalities. Physiologically Based Pharmacokinetic (PBPK) modeling remains a cornerstone for understanding antibody disposition across tissues and species^19–23^. While preliminary models for mRNA–LNP dynamics have begun to emerge^24,25^, they remain limited in scope and lack integration with systemic pharmacokinetic models, reducing their versatility for broader therapeutic applications. Even our previous work, Fiandaca *et al*. (2025)^26^, which well describes the trafficking from mRNA injection to blood samples, was validated on a single mRNA formulation and antibody, and it does not account for neonatal Fc receptor (FcRn) binding.

To address this gap, we extended the PBPK model from Sepp *et al*. (2019)^27^ with a novel LNP–mRNA layer to support the trafficking of mRNA-encoded therapeutics in mice. The resulting multiscale model was trained and validated on five preclinical datasets^28–32^, covering different antibody formats (with molecular weight ranging from 55kDa to 150kDa, and with or without the Fc region) and LNP formulations, and it incorporates:

- a phenomenological representation of LNP trafficking, cellular uptake, endosomal processing, and translation of mRNA into the encoded antibody, and
- an established PBPK framework for antibody disposition, incorporating FcRn binding and the two-pore hypothesis, and adapted to accommodate a broader range of mRNA-encoded therapeutics.

Overall, this integrative approach establishes a foundation for modeling of mRNA-based antibody therapeutics, offering a framework to explore delivery efficiency, expression kinetics, and systemic distribution across preclinical settings.

## Results

Our work aims to develop a flexible, robust, and general-purpose multiscale model capable of accurately predicting the pharmacokinetic (PK) dynamics of therapeutic antibody concentrations following administration of IVT mRNA formulations in mice.

In the first Section, we present the model from Sepp *et al*. (2019)^27^ and analyze its parameters, identifying the most sensitive terms to further generalize the model and enhance its robustness across different therapeutics.

In the second Section, we introduced a novel LNP–mRNA layer to phenomenologically describe the biological processes that span from the LNP–mRNA injection to the intracellular translation of the encoded antibody. With a minimal number of additional parameters, this layer was integrated into the existing PBPK structure to obtain a multiscale model for mRNA-encoded therapeutics.

Finally, we calibrated and validated the model against five independent datasets, demonstrating its reliability and versatility across a wide range of therapeutic antibodies, mRNA constructs, and LNP formulations.

### Therapeutic trafficking model and analysis of key parameters

The antibody trafficking module of our model was developed following the PBPK framework proposed by Sepp *et al*. (2019)^27^ and comprises 14 tissues, a *Blood* compartment, an additional *Tumor* compartment for oncology applications, and a *Other* compartment accounting for the remaining 2% of body volume and blood flow, as shown in Fig. 1. Each tissue is divided into vascular, interstitial, and endosomal sub-compartments. The uptake of Abs is governed by the two-pore hypothesis introduced in Rippe *et al*. (1994)^33^, which accounts for size-dependent transcapillary exchange.

**Figure 1:**
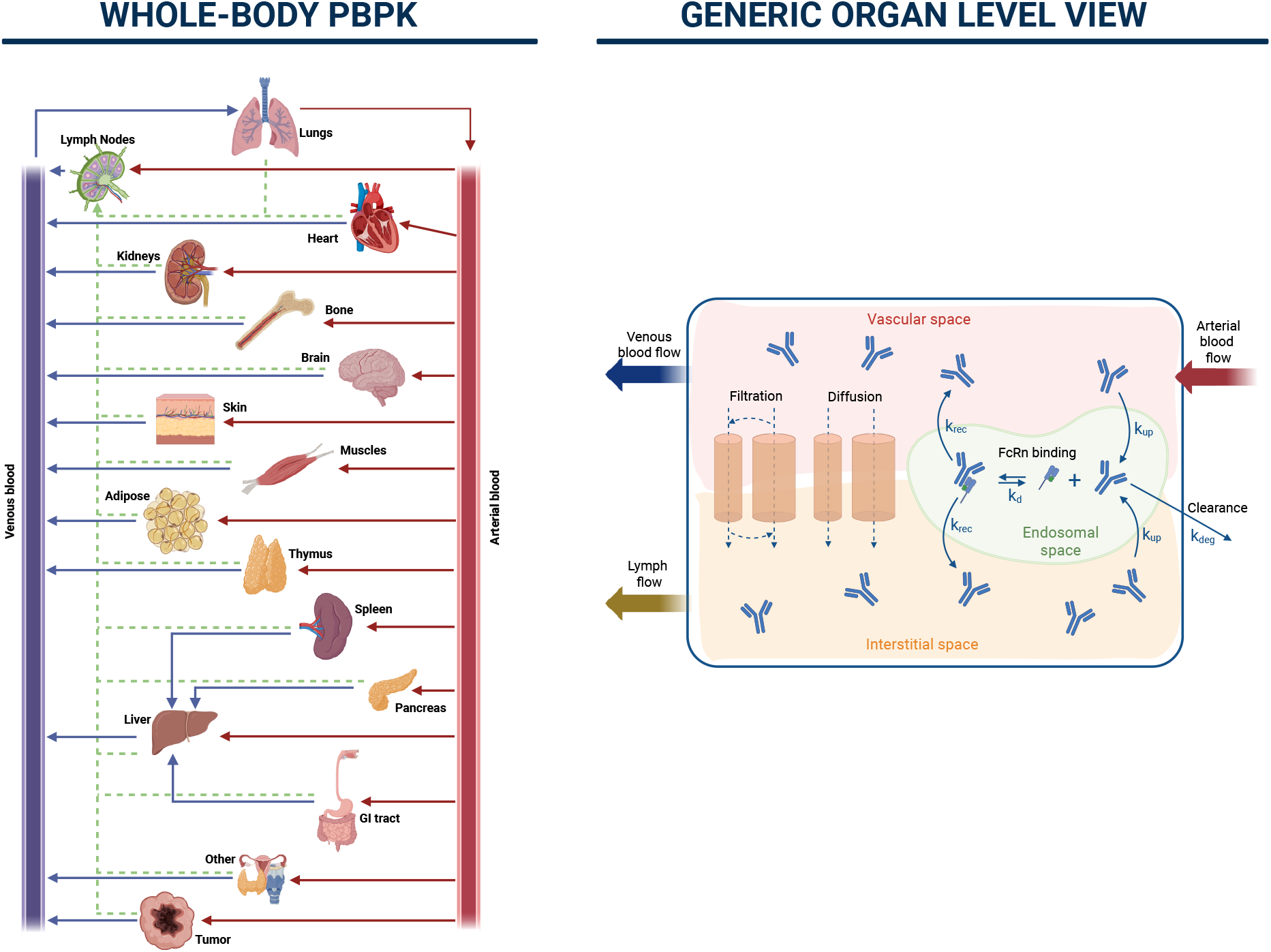
Representation of the PBPK model. The left panel illustrates the whole-body organization, where each compartment represents a specific tissue or organ connected by arterial, venous, and lymphatic blood flows. The right panel shows the generic organ-level structure, including vascular, interstitial, and endosomal spaces, and key mechanisms such as filtration, diffusion, FcRn binding, and clearance. Created in BioRender. Marchetti, L. (2026) https://BioRender.com/9lgli1o.

The PBPK model by Sepp *et al*. (2019)^27^ was selected as the foundation for this work because it provides a unified, two-pore platform capable of describing the disposition of proteins of different sizes, from small domain antibodies to full IgG molecules. Unlike earlier models that treated lymphatic flow and permeability parameters independently^19,22,34^, Sepp *et al*. introduced a mechanistically consistent relationship between organ-specific lymph flow and permeability–surface area products, reducing overparameterization and improving physiological realism. The model integrates endosomal FcRn-mediated recycling, allowing the simulation of both antibodies with and without Fc regions, and was evaluated across mice and rats using extensive tissue-distribution datasets. Moreover, Sepp *et al*. compared multiple physiological parameter sets and identified those that best captured tissue-specific kinetics, ensuring robust cross-species applicability.

Before modifying the model structure by introducing the LNP–mRNA layer, we first ensured that the PBPK framework remained versatile across diverse therapeutic products. To this end, we reviewed all drug-dependent parameters of the original PBPK model—listed below and shown in Fig. 1—and subsequently performed a global sensitivity analysis, informed by literature-based ranges, to determine which of them exerted the strongest influence on the model output.

- Molecular weight (MW) of the protein, used to compute permeability-related parameters via the two-pore model.
- Protein degradation rate (*k*_deg_), controlling the main elimination pathway occurring in the endosomal space of each tissue.
- Non-specific protein pinocytotic uptake rate (*k*_up_), determining transport from the vascular to the endosomal space in each tissue.
- FcRn binding and recycling rate constants (*k*_d_, *k*_rec_). These parameters define the fraction of antibody that binds FcRn and is recycled back into circulation; both are set to zero for proteins lacking Fc-binding domains.
- Scaling factor adjusting the effective interstitial space across tissues (*I*_adj_), accounting for steric and electrostatic effects related to antibody size and charge.

The two-pore model automatically adjusts permeability-related parameters based on the MW.

We computed a global Sobol’ sensitivity analysis^35^ on the remaining parameters to test their impact on the PBPK model output using Sepp *et al*. (2019)^27^ blood time series results (Fig. 2). The pinocytosis parameter *k*_up_, with high first- and total-order Sobol’ indices, exhibited by far the strongest impact on the model dynamics. Among the remaining parameters, *k*_d_ was the only one showing a noticeable contribution to the model output, with its effect arising mainly through parameter interactions rather than first-order effects. It is worth noting that *k*_d_ and *k*_rec_ are both directly related to FcRn binding and, together with *k*_deg_, were considered *a priori* to be irrelevant when the Fc domain is absent, based on the established FcRn-mediated recycling and protection from degradation of Fc-containing antibodies^20,34,36^. Based on these findings, *k*_deg_ and *k*_rec_ were fixed, whereas *k*_up_ and *k*_d_—the most protein-specific parameters—were retained as adjustable parameters during calibration. Several studies emphasize the importance of *k*_up_ (sometimes named *CL*_up_) and *k*_d_ (or their equivalents) in shaping PK profiles^23,34,37–40^. Both values need to take into account protein- and subject-dependent properties, such as changes in pH for FcRn binding and molecular charge for the pinocytosis. Since our model aims to capture a wide range of Abs, both parameters were fitted using broad intervals derived from the minimum and maximum values reported in the literature (see Materials and Methods).

**Figure 2:**
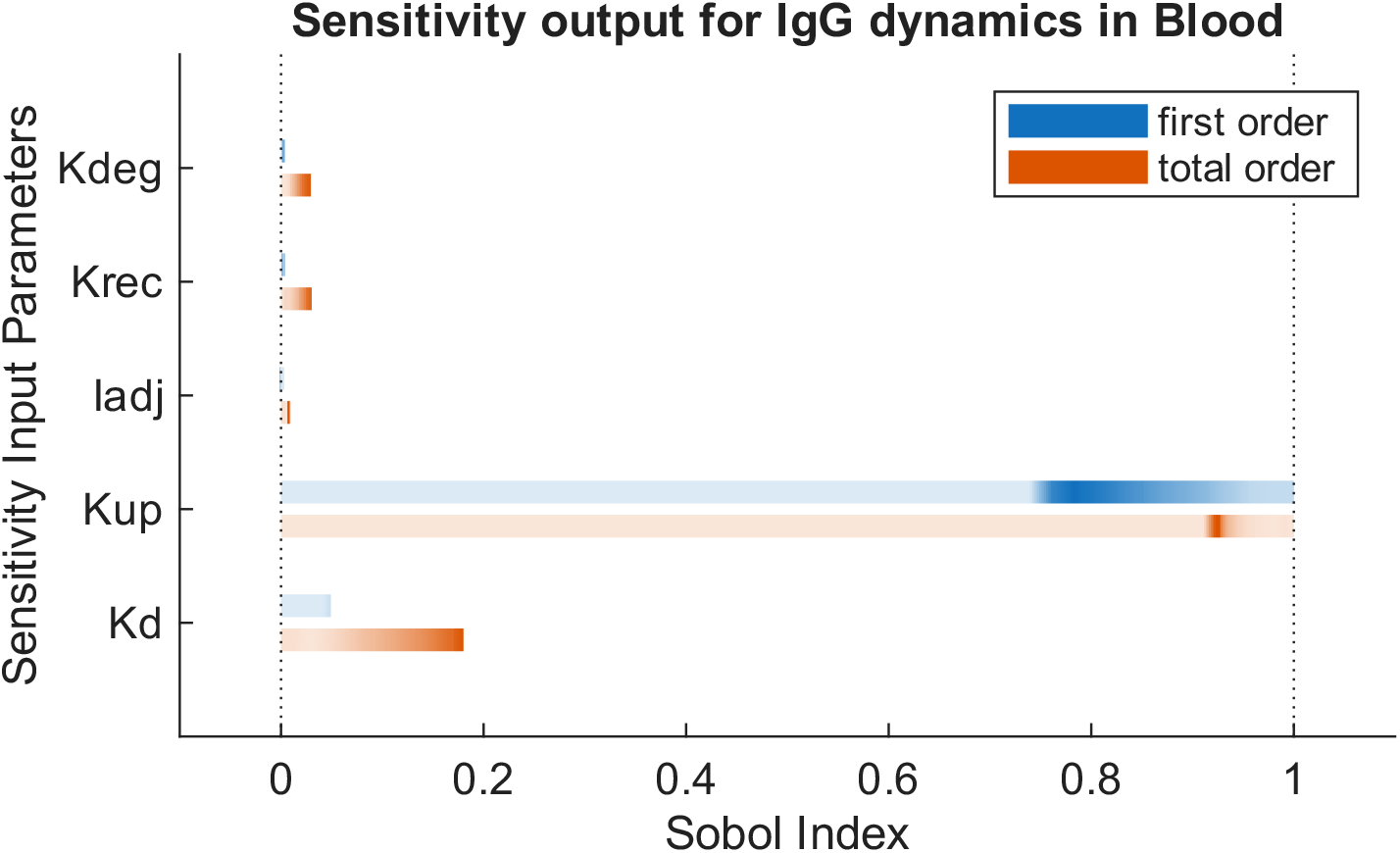
Sensitivity analysis results. Sobol’ sensitivity indices for the PBPK model parameters obtained using Sepp *et al*. (2019) blood time series as output. The plot reports first-order (blue) and total-order (orange) Sobol’ indices for each parameter, quantifying their relative contribution to the variance of the simulated blood concentration. Darker colors mean that those values occur more often over the whole time course.

The adjustment of interstitial space *I*_adj_ was first proposed by Wiig *et al*. (2008)^41^ and later adopted by Sepp *et al*. (2019), who showed that protein size and charge can reduce the effective interstitial space by up to 50%, due to steric and electrostatic effects within the interstitium. Since Sepp *et al*. reported modest AIC improvements (a relative variation of 0.2%) from halving interstitial volumes, we found this modification unjustified as a general rule. However, additional PK simulations using data from Li&Shah (2019) (see Figure S1 in Supplemental Material) showed improved fits for some larger proteins when this adjustment was applied^20^. Based on these findings and on the lack of knowledge regarding most antibody charges, we introduced a calibration factor (*I*_adj_ ∈ [0.5, 1]), following the input from Sepp *et al*., for recombinant proteins with molecular weights ≥ 100 kDa.

### Integration of an LNP–mRNA trafficking and translation layer

Our original contribution to the PBPK model structure is the integration of a phenomenological layer that describes all the processes occurring from LNP–mRNA administration to antibody synthesis. These include LNP-encapsulated mRNA trafficking, hepatic uptake, translation, and degradation. A schematic representation of this new model component is shown in Fig. 3. The additional layer consists of three main compartments: *i*) *Blood, ii*) *Hepatocyte Cells* within the interstitial space of the liver, and *iii*) a lumped *Other Organs* compartment encompassing all non-hepatic tissues. The mathematical formulation of this layer is reported in Eqs. 1–6, and its structure is described in the following.

**Figure 3:**
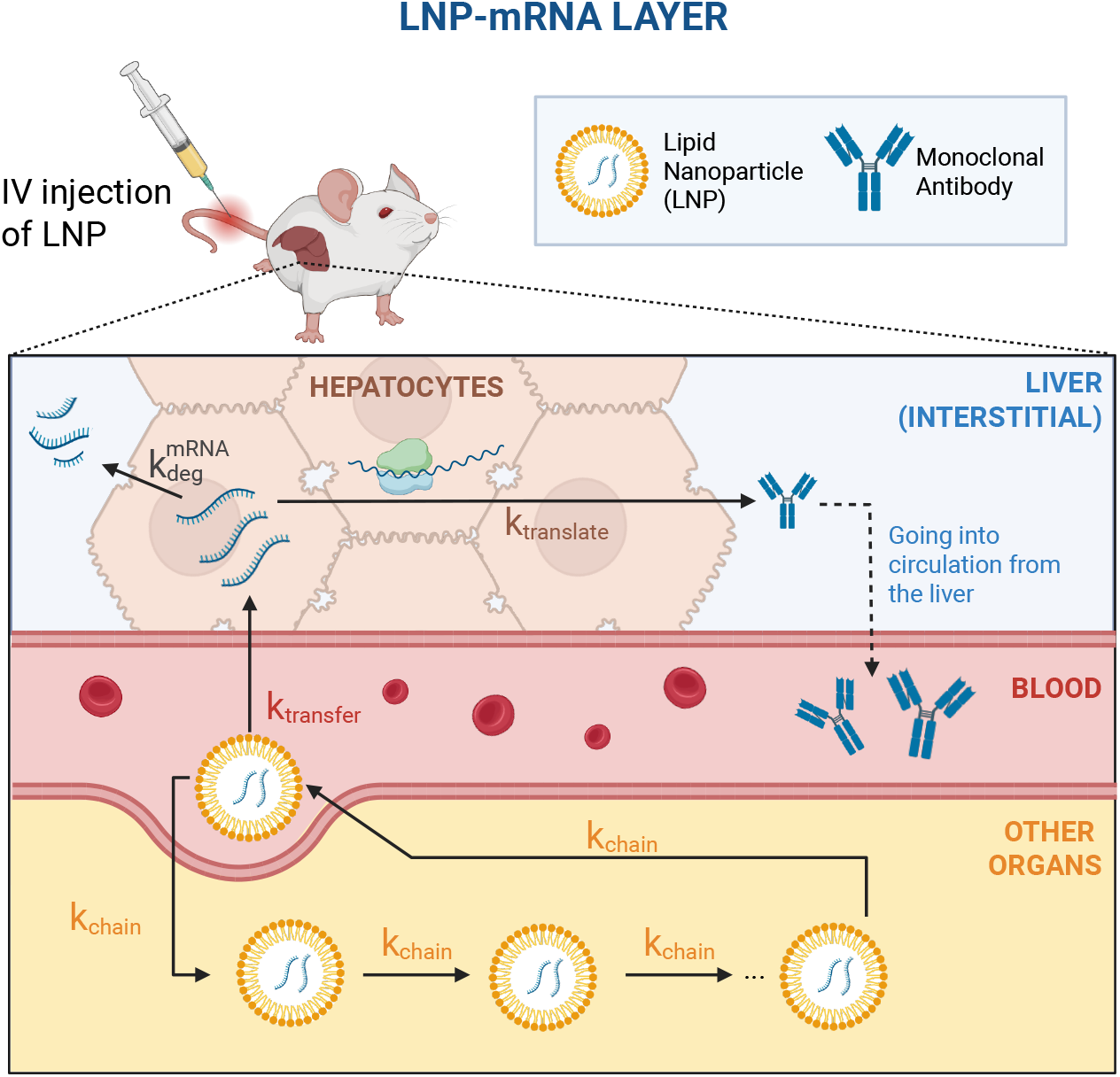
Representation of the novel LNP–mRNA layer. Schematic representation of the novel LNP–mRNA layer integrated into the PBPK model. The layer captures the trafficking of lipid nanoparticles (LNPs), intracellular mRNA release, translation into encoded antibodies, and subsequent secretion into systemic circulation. Created in BioRender. Marchetti, L. (2026) https://BioRender.com/2esof9m.

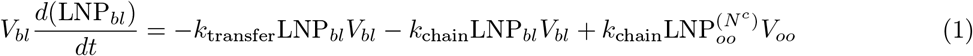

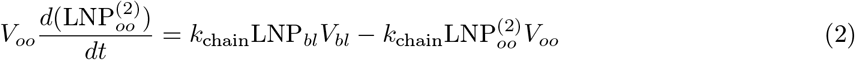

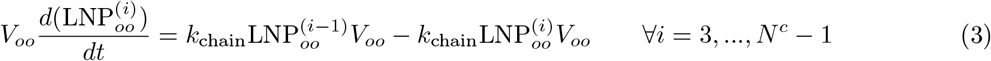

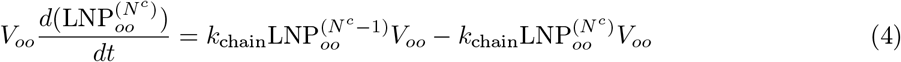

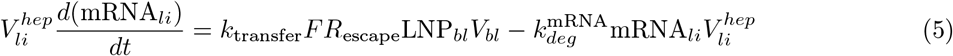

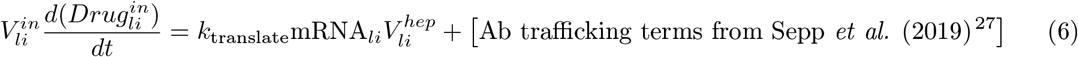

After intravenous injection, LNP-encapsulated mRNA enters the bloodstream. From there, it can circulate throughout the *Other Organs* or be taken up by the liver. The circulation throughout the *Other Organs* is modeled using a chain of compartments (Eqs. 2–4). Fluxes among the chain compartments are modeled as first-order reactions, parameterized by a common rate *k*_chain_. The number of compartments (*N*^*c*^) is variable and reflects the complexity and variability of LNP trafficking through the body. This trafficking process was introduced to mimic the biphasic profile consistent with both experimental data and literature reports^14^. By approximating a delay differential equation, our transit compartment model reproduces this behavior as a simplified alternative to a full PBPK model for LNPs. The goal is to avoid unnecessarily complex model structures by selecting, for each drug, the minimum number of compartments that adequately capture the observed data (details on the implementation in Materials and Methods).

Importantly, this compartment chain is not intended to represent re-entry of internalized LNPs into the systemic circulation as intact particles. Rather, it provides a phenomenological description of vascular transit and delayed distribution, including repeated passage through the circulation and reversible distribution between blood and tissue-associated spaces prior to irreversible cellular uptake. This simplification is motivated by the fact that early LNP pharmacokinetics are strongly influenced by formulation-dependent properties, particularly PEG-lipid composition and desorption behavior, which affect circulation time and biodistribution^42,43^.

During this process, a fraction of LNPs is redirected to the liver at each passage, thereby ensuring progressive accumulation in hepatocytes. This second pathway is governed by the rate *k*_transfer_ and accounts for both the transfer of LNPs to the liver and the subsequent endosomal escape of mRNA (Eq. 5). The parameter *FR*_escape_ defines the fraction of mRNA successfully released into the cytosol, representing the endosomal escape bottleneck that remains a major challenge in mRNA–LNP therapeutics^44^.

Once released, mRNA degrades with a first-order rate constant 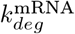 (fixed from the literature, see Table 1), while the remaining transcripts are translated into the therapeutic antibody through a reaction that combines translation and exocytosis into a single term governed by *k*_translate_ (Eq. 6). The resulting proteins enter systemic circulation, beginning from the liver interstitial space, and follow PBPK-based transport equations throughout the body.

**Table 1:** Parameters used in Eq. 1–6 of the mRNA-LNP layer. Fixed physiological values were obtained from literature, while fitted parameters were estimated from experimental data as reported in Table 2. When applicable, corresponding formulae and data sources are indicated.

| Parameter | Description | Value(Unit) | Formula | Source |
| --- | --- | --- | --- | --- |
| $V_{bl}$ | Blood volume | 0.379 (mL) | - | Sepp *et al.* (2019)<sup>27</sup> |
| $V_{oo}$ | "Other organs" volume | 25.68 (mL) | $\sum_{org \neq \text{liver, blood}} V_{org}$ | Shah *et al.* (2012)<sup>22</sup> |
| $V_{li}^{\text{in}}$ | Liver interstitial volume | 0.385 (mL) | - | Sepp *et al.* (2019)<sup>27</sup> |
| $V_{li}^{\text{hep}}$ | Hepatocyte volume | 0.861 (mL) | $0.7 \cdot V_{\text{liver}}^{\text{cell}}$ | Shah *et al.* (2012),<br>Sepp *et al.* (2019)<sup>22,45</sup> |
| $FR_{\text{escape}}$ | Fraction of LNP reaching the cytosol | 0.02 (-) | - | Liu *et al.* (2024)<sup>44</sup> |
| $k_{\text{deg}}^{\text{mRNA}}$ | mRNA degradation rate in liver | 0.0693 (1/h) | $\ln(2)/T_{\text{deg}}$ | Yang *et al.* (2003)<sup>46</sup> |
| $N^c$ | Number of transit compartments in Other Organs | Fitted (-) | - | Table 3 |
| $k_{\text{chain}}$ | Transit rate for the chain of compartments | Fitted (1/h) | - | Table 3 |
| $k_{\text{transfer}}$ | LNP transfer rate from Blood to Hepatocyte Cells | Fitted (1/h) | - | Table 3 |
| $k_{\text{translate}}$ | Efficacy rate for mRNA-encoded Ab translation | Fitted (1/h) | - | Table 3 |

In the Eqs. 1–6, LNP_*bl*_, 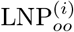, mRNA_*li*_, and 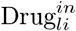 represent, respectively, the LNP concentrations in the blood and in the *i*-th compartment of Other Organs, the mRNA concentration in the liver, and the drug concentration in the liver. The constant parameters from the equations are summarized in Table 1 together with their descriptions, values, and literature references.

### Calibration and validation across multiple mRNA-encoded antibody datasets

The model was calibrated and validated using datasets describing the concentration-time profiles of five mRNA-encoded antibodies: B7H3×CD3 Bispecific T-cell Engager, RiboMab02.1, XA-1, Pembrolizumab, and Trastuzumab (see Table 2).

**Table 2:**
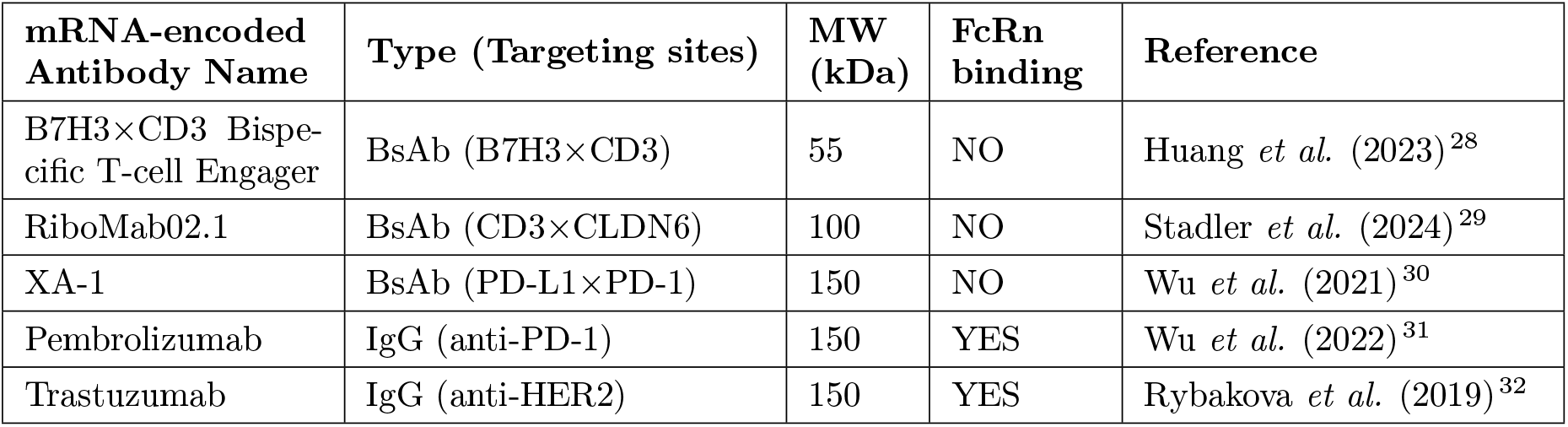
Dataset and molecular properties of mRNA-encoded antibodies. Summary of mRNA-encoded therapeutic antibodies used for model training and validation, including their molecular characteristics and FcRn binding status.

For each molecule, the model was fitted using both recombinant and mRNA-derived data, yielding simulated profiles that closely reproduced the observed time courses (Fig. 4). The two-step training procedure described in Materials and Methods was applied independently to all datasets, yielding consistent parameter estimates across therapeutics (Table 3). Given the phenomenological nature of the new layer and the limited knowledge of LNP–mRNA movements and reactions, precise ranges for the parameters *k*_chain_, *k*_transfer_, and *k*_translate_ are not yet established. For this reason, an identifiability analysis was performed prior to calibration on the mRNA-related parameters as outlined in Materials and Methods, resulting in local structural identifiability. This result should nevertheless be interpreted with caution, as it applies only locally to the phenomenological layer. Since the analysis was performed without imposing bounded parameter ranges, additional data or more specific biological constraints may further refine the parameterization and improve its biological interpretation. To further assess parameter robustness under the available data conditions, 95% confidence intervals for the fitted parameters are reported in Supplemental Table S1. By separating the training process, we minimized dependencies between drug-specific and mRNA-specific parameters.

**Table 3:** Fitted parameters for each dataset. The parameters were estimated individually for each study based on single-dose plasma kinetics. Boxes marked with “–” indicate that the parameter did not require fitting for that drug. When not fitted, *I*_adj_ is set to 1 and *K*_*d*_ to 0 M.

| Parameter (Unit) | Huang2023<sup>28</sup> | Stadler2024<sup>29</sup> | Wu2021<sup>30</sup> | Wu2022<sup>31</sup> | Rybakova2019<sup>32</sup> |
| --- | --- | --- | --- | --- | --- |
| $k_{up}$ (1/h) | 11.88 | 10.46 | 0.1107 | 0.2193 | 0.07104 |
| $k_d$ (M) | - | - | - | $6.978 \times 10^{-6}$ | $4.771 \times 10^{-5}$ |
| $I_{adj}$ (-) | - | - | 0.5 | 1 | 0.5 |
| $N^c$ (-) | 6 | 2 | 4 | 4 | 6 |
| $k_{chain}$ (1/h) | 0.2305 | 0.03985 | 0.01937 | 0.02808 | 0.08446 |
| $k_{transfer}$ (1/h) | 0.1257 | 0.1876 | 0.01390 | 0.03421 | 0.09494 |
| $k_{translate}$ (1/h) | 106.22 | 534.6 | 61.30 | 52.38 | 23.02 |

**Figure 4:**
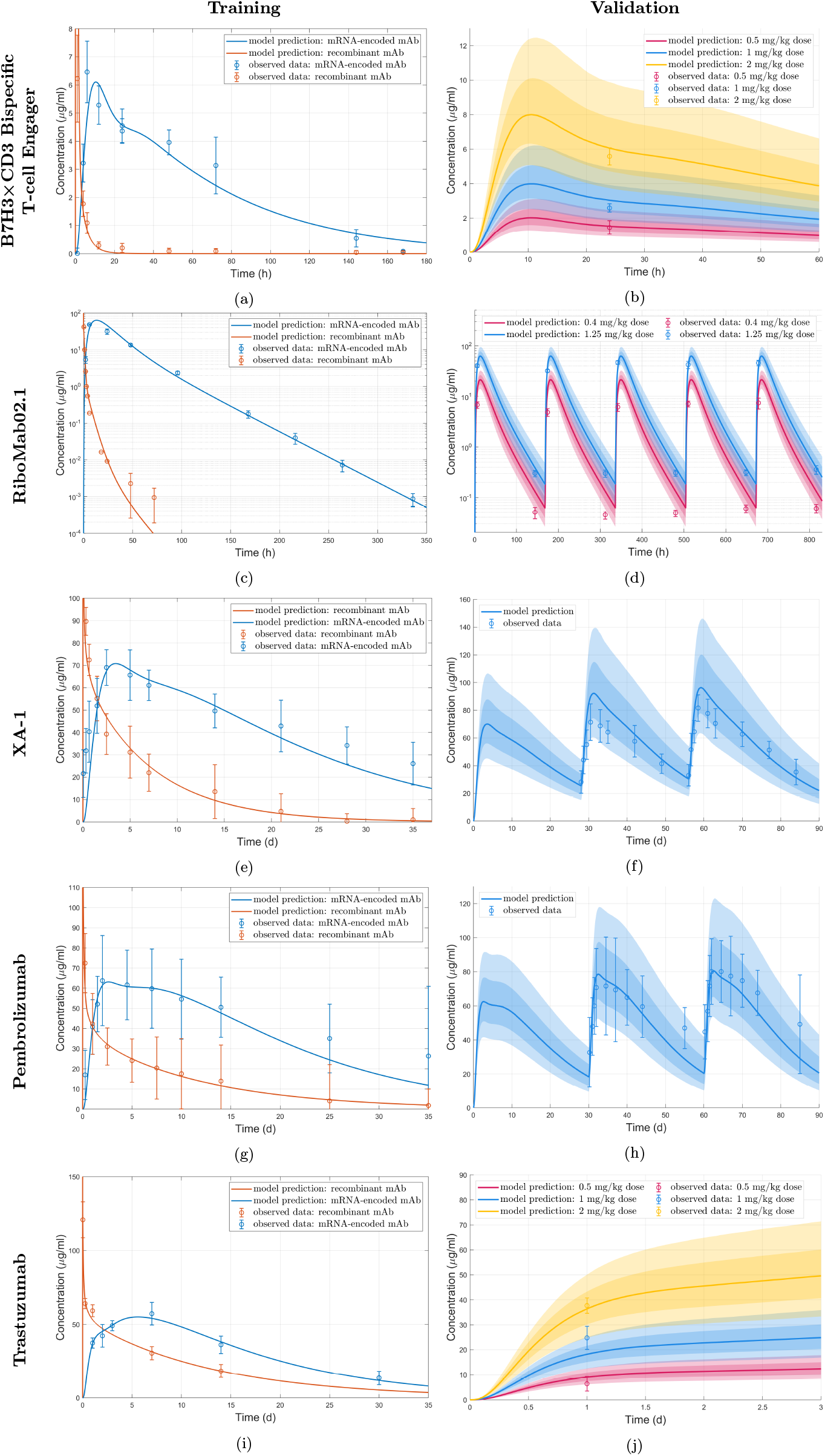
Calibration and validation results for the five therapeutics. The names in vertical refer to the five therapeutics described in Table 2. Left panels show calibration time series for recombinant antibodies (orange curves) and their mRNA-encoded counterparts (blue curves), using single-dose PK data. Right panels show validation results for mRNA-encoded antibodies, where shaded bands represent the 10th, 25th, 75th, and 90th percentiles generated through Monte Carlo simulations. All dosages are intravenously injected. Training doses (left panels): (a) Recombinant: 6 mg/kg, mRNA-encoded: 1.5 mg/kg mRNA (LNP); (c) Recombinant: 2.5 mg/kg, mRNA-encoded: 1.25 mg/kg mRNA (LNP); (e) Recombinant: 10 mg/kg, mRNA-encoded: 2 mg/kg mRNA (LNP); (g) Recombinant: 10 mg/kg, mRNA-encoded: 2 mg/kg mRNA (LNP); (i) Recombinant: 8 mg/kg, mRNA-encoded: 2 mg/kg mRNA (LNP). Validation doses (right panels): (b) 0.5 mg/kg, 1 mg/kg, and 2 mg/kg (single administrations); (d) 0.4 mg/kg and 1.25 mg/kg (five-dose regimen, every 7 days); (f) 2 mg/kg (three-dose regimen, every 28 days); (h) 2 mg/kg, (three-dose regimen, every 30 days); (j) 0.5 mg/kg, 1 mg/kg, 2 mg/kg (single administrations).

Model calibration was performed using single-dose data, whereas validation was executed against independent dose levels or multidose regimens. The time series of recombinant Abs and their mRNA-encoded counterparts (left panels of Fig. 4) showed clear and consistent kinetic differences, with the latter exhibiting prolonged circulation times and delayed peaks, in agreement with their expected intracellular production and secretion dynamics.

In the validation plots (right panels of Fig. 4), intersubject variability was represented through percentile bands derived from Monte Carlo simulations, where selected parameters were perturbed around their calibrated values (details in Materials and Methods). The shaded areas encompass most experimental observations, indicating that the estimated uncertainty adequately reflects the observed data dispersion.

The accuracy of the simulations in reproducing the experimental data was assessed using the mean absolute average fold error (AAFE), which quantifies the agreement between predicted and observed values (all values in Supplemental Table S2).

Overall, the AAFE across datasets was 1.44 for recombinant antibody fitting, 1.36 for mRNA-encoded fitting, and 1.27 for validation, indicating deviations typically below twofold. Across mRNA-encoded antibody calibrations, residuals were within or near the experimental uncertainty, and Bayesian Information Criterion (BIC) values confirmed that the model achieved an appropriate trade-off between model complexity and fit quality.

## Discussion

In this work, we developed an integrative PBPK modeling framework for mRNA-encoded therapeutic antibodies in mice. The model builds on the well-established antibody trafficking model by Sepp *et al*. (2019)^27^ and extends it with a phenomenological LNP–mRNA trafficking layer describing its uptake, intracellular translation, and degradation. The physiological structure of the original PBPK model to describe the antibody circulation was retained, while key drug-specific parameters (*k*_up_, *k*_d_, *I*_adj_) were re-estimated for each antibody alongside the parameters of the phenomenological layer describing the LNP–mRNA trafficking (*k*_chain_, *k*_transfer_, *k*_translate_). The adopted two-step calibration approach relies on the assumption that the recombinant and mRNA-encoded versions of each therapeutic are either identical or differ only in ways that are negligible for protein trafficking. The parsimonious multiscale framework was calibrated and validated on five datasets spanning recombinant and mRNA-encoded antibodies with diverse molecular weights, Fc properties, and LNP formulations, demonstrating robust predictive performance.

Our group recently developed an mRNA trafficking model^26^ coupled with the PBPK model introduced by Li&Shah (2019)^20^. For the present work, we instead adopted Sepp *et al*. (2019)’s PBPK model, as it incorporates antibody recycling via FcRn binding, thereby enabling more accurate modeling of Fc-containing Abs and supporting the use of a broader range of datasets for model training and validation. Furthermore, our previous effort^26^ assumed direct hepatic delivery of LNPs, while the current model emphasizes extravasation and redistribution prior to liver uptake, drawing inspiration from data analysis and the insights provided by Zhou *et al*. (2025)^14^. To describe such biological insights, we introduced a chain-compartment model in which both the number of compartments (*N*^*c*^) and the rate constant (*k*_chain_) determine the effective delay associated with LNP movement through peripheral tissues before re-entering the plasma circulation. To further support the need for a circulation-related delay in the model, we fitted the mRNA-related parameters under the assumption of no transit compartment, i.e., by setting *k*_chain_ = 0 *h*^*−*1^. The resulting fits, shown in Figure S2 of the Supplemental Materials, clearly demonstrate that the model fails to capture the pharmacokinetics of the mRNA-encoded therapeutic in the absence of this delay mechanism. Although the transit-compartment chain improves the description of the early biphasic profiles, its interpretation remains phenomenological. In the present framework, *k*_chain_ and *N*^*c*^ should not be interpreted as direct surrogates of specific anatomical pathways, but rather as effective descriptors of delayed LNP redistribution prior to irreversible uptake. Since varying *N*^*c*^ modifies the model structure itself, its identifiability is intrinsically more difficult to formalize than that of continuous parameters. Nevertheless, under the adopted *BIC*^*new*^-based selection procedure (details in Materials and Methods), each product showed a well-defined optimum, supporting the practical usefulness of the selected *N*^*c*^ values within the current phenomenological framework.

The variation of *k*_chain_, *k*_transfer_, and *k*_translate_ across products likely reflects formulation-dependent differences in delivery and expression behavior. In the present framework, all three parameters should be interpreted as lumped effective descriptors rather than as direct physicochemical correlates of specific LNP properties. More specifically, *k*_chain_ captures early redistribution and circulation delay, *k*_transfer_ captures productive liver-directed delivery, and *k*_translate_ captures downstream expression processes, including translation and secretion.

Despite recent efforts to develop PBPK models focused on LNP trafficking^24,25^, we believe that, given the current state of experimental data and biological understanding, a more parsimonious modeling strategy may be better suited to reliably capture mRNA-encoded therapeutic pharmacokinetics. At present, the available data and biological knowledge are still insufficient to robustly support highly detailed model representations involving numerous uncertain parameters, such as organ-specific LNP uptake rates. Moreover, the available literature data are largely restricted to plasma concentration–time profiles, with little or no tissue-resolved information on intracellular mRNA kinetics or LNP biodistribution. Such measurements would be essential to support more mechanistic and better-constrained model formulations in future studies. In this context, a key strength of our approach is its ability to reproduce diverse datasets using a minimal and effective structure, thereby limiting overfitting and reducing reliance on poorly constrained assumptions.

In the present model, *FR*_escape_ and 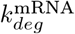 were fixed across datasets to limit the degrees of freedom introduced by the novel mRNA-related layer, given the limited data currently available. We acknowledge that this is a simplification, as both processes may depend on formulation-specific properties of the mRNA–LNP product. To reduce the uncertainty associated with this assumption, we restricted the analysis to datasets based on relatively similar conventional LNP formulations, and 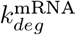 was included among the perturbed parameters used to generate the variability bands shown in the validation plots. More informative measurements of LNP and intracellular mRNA dynamics will be needed to estimate these parameters in a product-specific manner in future model extensions. At the same time, this phenomenological representation may limit the direct translatability of the model across species and could necessitate structural refinements to achieve reliable predictions in clinical settings. Our group is currently working on a more mechanistic—even though still parsimonious in nature—translational layer, leveraging qualitative information about the LNP’s fate and adapting it to different preclinical animal species.

In the present framework, hepatocytes were considered the dominant site of antibody production, consistent with the biology of conventional liver-tropic LNPs. For standard ionizable formulations, ApoE-mediated uptake and the hepatic microenvironment favor functional delivery to hepatocytes, whereas uptake by other hepatic or extrahepatic cell types does not necessarily translate into equally relevant protein secretion. Accordingly, for the datasets analyzed here, we considered a hepatocyte-driven production term to be the most biologically appropriate representation^47–49^. Future extensions may nevertheless evaluate explicit non-hepatic production compartments when tissue-resolved expression data become available or when considering formulations with broader tropism.

Despite our efforts to make the Sepp *et al*. (2019) model versatile for antibodies of different sizes, we observed a loss of accuracy in reproducing the terminal data points for the smaller recombinant therapeutics (Fig. 4a, 4c). The two curves yielded AAFE values of 1.64 and 1.87, respectively—still adequate but higher than the average value of 1.2 observed for the other three recombinant therapeutics. This behavior also corresponds to the estimated *k*_up_ ~10 *h*^*−*1^ in both cases (Table 3), which, while within plausible limits, remains higher than the average reported in the literature.

In the case of the B7H3×CD3 Bispecific T-cell Engager, a relatively high AAFE of 1.78 was also obtained for the mRNA-encoded antibody (Fig. 4a), likely due to the accentuated biphasic behavior observed in the data. The same trend is reflected in the fitted parameter values (Table 3), where both the number of compartments *N*^*c*^ and the rate *k*_chain_ are elevated. This slight loss of accuracy results from our effort to balance model complexity and predictive performance, as quantified by the modified Bayesian information criterion *BIC*^*new*^, which led to the selection of a shorter compartment chain (details in Materials and Methods).

For the remaining mRNA-encoded therapeutics, the model accurately reproduced the dynamics: RiboMab02.1 (Fig. 4c) and Trastuzumab (Fig. 4i) achieved AAFEs of 1.15 and 1.06, respectively, while XA-1 (Fig. 4e) and Pembrolizumab (Fig. 4g) maintained AAFEs of 1.42 and 1.37. The latter two values can still be considered highly accurate, given the relatively high experimental uncertainty—particularly for Pembrolizumab—and despite the statistical measure not explicitly accounting for it, all fitted simulations remained within or near the experimental standard deviation.

The estimate of the mRNA translation efficacy *k*_translate_ might seem high, but it is well with the values suggested by previous PK models^24,25^.

During validation, all therapeutics—except RiboMab02.1 (Fig. 4d)—achieved validation AAFE values between 1.1 and 1.25.

The case of RiboMab02.1, together with Fig. 4b and 4j, illustrates that the model’s predictive performance can still improve under dosage variations.

The XA-1 and Pembrolizumab validation plots (Fig. 4f and 4h), both corresponding to multidose regimens, reveal distinct behaviors in the response to repeated administrations. Although the model reproduces both datasets well, it slightly overestimates the experimental data in Fig. 4f and underestimates them in Fig. 4h, suggesting possible differences in immunogenic response related to the product, such as accelerated blood clearance (ABC) observed for several lipid-based delivery vehicles, the experimental subjects, or both^50–52^. From a modeling perspective, this phenomenon could be represented through an additional component regulated by anti-PEG antibodies^53–55^. Such an extension could improve predictive performance in multidose regimens by allowing effective clearance to vary across repeated administrations, thereby capturing immune-driven nonlinearities that are beyond the scope of a purely linear framework. Since the datasets considered here do not include direct measurements of anti-PEG antibodies or related immune biomarkers, we leave ABC-based formulations as an important future extension of the model.

Exploring the many papers and datasets available in the literature, we carefully selected those that fulfilled key criteria: a well-sampled and consistent time series for both recombinant and mRNA-encoded therapeutics, and a description of critical drug and mRNA–LNP properties (see Table 2).

In addition to the adopted datasets, two further studies were considered^56,57^. Despite their promising results, the study by Stadler *et al*. (2017) employed a lipid-based formulation that differs from conventional LNPs and reported only a single time series, limiting its applicability. Meanwhile, the study by Bähr-Mahmud *et al*. (2023), while demonstrating stable systemic exposure in animal models, provided pharmacokinetic data with limited temporal resolution and sample size, reporting only a few plasma time points with group-average concentrations. Consequently, despite the significance of their findings, these limitations precluded the robust parameter identification required for PBPK model calibration, and their dataset was therefore respectfully excluded from our analysis. It is also important to note that the data in Wu *et al*. (2022)^31^ includes extensive datasets—covering both single- and multi-dose treatments—but individual data points often exhibit considerable variability, and some inconsistencies are evident across plots. We found it necessary to exclude one dosage-related plot (Fig. 2A in Wu *et al*. (2022)^31^) because the initial value at the highest dose was nearly three times higher than in the following two plots (Fig. 2B and Fig. 3 in Wu *et al*. (2022)^31^), despite identical dosage and timepoint, raising concerns about data reliability.

Even with this stringent selection, challenges emerged when validating dose–response curves. As noted above, datasets showed slight over- or under-predictions, which may reflect variability in mice or could suggest saturation effects and immune-driven responses, often without a consistent underlying pattern. One experiment from Wu *et al*. (2021)^30^ was also excluded from the validation, as it displayed pronounced non-linear variations across doses that could not be captured within the current model formulation. Nevertheless, despite its deliberately simple structure, our model was able to reproduce observations across substantial ranges of therapeutics (see validation plots: Fig. 4, right panels).

Rather than constituting model failures, these discrepancies highlight important limitations of the present framework and point to the need to investigate and integrate more detailed mechanisms of immune activation and dose-dependent non-linearities, which could ultimately enhance the predictive power of PBPK approaches for mRNA-encoded therapeutics. Thus, while some datasets challenge the current framework, they also provide valuable guidance for the next generation of models.

Our review of the available literature also revealed that most mRNA-encoded therapeutic studies to date have focused on oncology applications, with only a few examples targeting other diseases, such as Chikungunya virus infection^58,59^. Although the structure of our model is broadly applicable and could, in principle, be adapted to non-oncological contexts, the predominant availability of cancer-related data guided our decision to focus the present analysis and simulations on mRNA-encoded therapeutics for cancer. Nonetheless, extending this framework to additional disease contexts remains an active area of work in our group, supported by the increasing availability of diverse experimental data^60^.

Future developments will aim to take advantage of the rapidly expanding amount of experimental data on mRNA–LNP therapeutics to progressively refine the present framework. In particular, our group is working toward the integration of more mechanistic representations of intracellular processes and immune responses, potentially through Pharmacodynamic or Quantitative Systems Pharmacology extensions. These additions would enable the model to capture product-specific mechanisms, such as target engagement, immune activation, and translation efficiency, while maintaining the physiological interpretability of the PBPK structure. Progress toward a more mechanistic description of LNP trafficking will depend on the generation of shared experimental measurements that directly relate formulation features to biological observables. In future model extensions, this may enable replacement or integration of the current transit-compartment representation with explicit physiological processes, such as organ-specific vascular exchange, lymphatic drainage, and tissue-specific uptake pathways. In the long term, such mechanistic enrichment may support the development of *ad hoc* models for different therapeutic products and ultimately pave the way for model-informed strategies toward personalized mRNA therapeutics, where dosing, formulation, and expression kinetics can be optimized on an individual basis. While the current framework was developed primarily for hepatic delivery of mRNA–LNP therapeutics, where hepatocytes act as the main sites of uptake and translation, alternative administration routes such as intramuscular (i.m.) and subcutaneous (s.c.) are increasingly relevant for clinical applications. These routes typically promote more localized effects and can prolong mRNA and LNP stability at the injection site, influencing both absorption kinetics and systemic exposure^13^. Future work will focus on extending the model to incorporate these administration routes, particularly intramuscular delivery, given that several i.m.-administered mRNA–LNP therapeutics have already received FDA approval^14^.

In parallel, the adopted PBPK framework—by retaining its mechanistic structure—preserves an intrin-sic ability for interspecies rescaling^22,27,61^. Ongoing efforts in our group aim to exploit this property to enable translation across preclinical species and eventually to humans, supporting the development of predictive tools capable of optimizing therapeutic kinetics and guiding the design of individualized mRNA treatment strategies.

In summary, this study establishes a PBPK framework grounded in physiological principles and extended with a parsimonious LNP–mRNA layer capable of capturing the key pharmacokinetic features of mRNA-encoded antibody therapeutics. The model was successfully validated across five independent antibodies, demonstrating its robustness and versatility in reproducing diverse pharmacokinetic behaviors. By integrating this minimal, phenomenological layer within an established physiological model, our work provides a flexible platform for quantitative exploration and design optimization of emerging mRNA-based therapies. As the field continues to evolve, such integrative modeling approaches will be instrumental in advancing model-informed development and accelerating the clinical translation of next-generation mRNA therapeutics.

## Materials and Methods

### Data

To develop our model, we used a broad set of publicly available pharmacokinetic data from the literature on mRNA-encoded therapeutics in mice. The five antibodies analyzed differ in several biological aspects, most notably in molecular weight and their affinity for the Fc receptor. Both features are crucial for understanding trafficking dynamics: molecular weight influences organ uptake, while the presence of the Fc region can significantly impact the protein’s half-life. In addition, mRNA and LNP formulations can affect the efficiency of antibody translation and influence aspects of trafficking. Therefore, it is important to characterize each therapeutic product and its key properties. A summary of the therapeutics used in this study, along with their relevant characteristics, is provided in Table 2. Data were extracted from the published studies either directly from tables or, when necessary, digitized from graphical representations, and then converted into consistent units before model calibration. Because the datasets originated from independent studies, differences in experimental design, sampling schedules, and bioanalytical methods may also contribute to inter-study variability.

All dosages were adjusted for a 28 g mouse, consistent with the physiological parameters used in the original PBPK model^27^. Specifically, for each experiment, we identified the weight *w* and dosage *d* and simulated the model using a dose normalized as follows: *d*^*normalized*^ = *d* · *w*_0_*/w*, where *w*_0_ = 28 *g*.

### Calibration and validation methods

Since each of the selected studies in the Data section provides PK data for both recombinant and mRNA-encoded versions of the same antibody, we performed model training in two sequential steps for each therapeutic. In the first step, we calibrated the drug-dependent parameters of the PBPK antibody layer using the recombinant antibody PK data. Specifically, we estimated *k*_up_ and, when required by the molecular properties of the therapeutic, also *k*_d_ and *I*_adj_. The parameters estimated for each dataset are summarized in Table 3, while boxes marked with “–” indicate parameters that were kept fixed. As already mentioned in Results, these parameter calibration ranges are derived from their minimum and maximum literature values, and, similarly for sensitivity analysis purposes only, we defined analogous ranges for the remaining therapeutic key parameters *k*_rec_ and *k*_deg_ (see Supplemental Tables S3–S5 for details).

In the second step, the parameters estimated from the recombinant antibody data were fixed, and the mRNA-encoded antibody PK data were used to calibrate the mRNA–LNP layer. More specifically, for each product we first estimated the mRNA-related parameters *k*_chain_, *k*_transfer_, and *k*_translate_ using at least one mRNA-encoded time series per therapeutic (see Supplemental Table S3 for details) and then determined the length of the transit-compartment chain, as further detailed in the next subsection. This sequential strategy was adopted to separate protein-specific PK features from mRNA–LNP-specific delivery and expression processes.

All validation plots display the inter-subject variability of the resulting time series as percentile bands. Variability was assessed through a Monte Carlo analysis comprising 10,000 simulations, each performed with perturbations of the following model parameters: *k*_up_, *k*_d_, *k*_deg_, 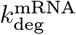, *k*_chain_, *k*_transfer_, and *k*_translate_.

For each simulation, parameter values were randomly sampled from normal distributions 𝒩 (*µ, σ*^2^), where *µ* corresponds to the parameter value and *σ* = 0.2 ∗*µ* (i.e., 20% of the mean). Simulations were then performed for all samples, and the median together with the 10th, 25th, 75th, and 90th percentiles were plotted for each scenario.

### Statistical measures

As mentioned in Results, the number of compartments may vary depending on both the subject and the administered LNP. Accordingly, *N*^*c*^ was determined in a data-driven manner through multiple calibrations against the mRNA-encoded time series. To this end, following similar approaches^62,63^, we employed a modified version of the Bayesian Information Criterion (BIC):

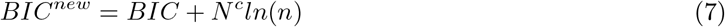

where *n* denotes the number of data points used for parameter estimation. Existing criteria in the literature typically compare models that differ in their number of parameters, thereby motivating the development of a new criterion for our application. In our framework, varying the chain length increases the number of compartments—and thus the effective degrees of freedom available during parameter calibration—while the total number of estimated parameters remains constant. Therefore, this statistical measure allows us to account, in a single quantity, for both goodness of fit through the *BIC* term and model complexity through the *N*^*c*^*ln*(*n*) penalty. Since *N*^*c*^ modifies the model structure, for each product we performed separate calibrations of the three mRNA-related parameters for discrete values *N*^*c*^ = 1, …, 15 and computed the corresponding *BIC* values. The optimal value of *N*^*c*^ was then selected as the one minimizing the modified criterion *BIC*^*new*^, i.e. 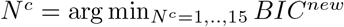. The resulting *BIC*^*new*^(*N*^*c*^) profiles for all products are reported in Supplementary Figure S3, showing a fairly convex trend across the tested chain lengths and a corresponding well-defined minimum for the selected value of *N*^*c*^.

To quantify the discrepancy between model predictions and observations in both calibration and validation, we computed the Absolute Average Fold Error (AAFE)^64^. The AAFE measures the average fold difference between predicted and observed values and is defined as:

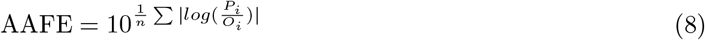

where *O*_*i*_ are the observed experimental values at the times *T*_*i*_, *P*_*i*_ are the corresponding model predictions, and *n* is the number of paired data points. By construction, *AAFE* ≥ 1, and values below 2 are generally considered to reflect good predictive performance^64^.

### Model implementation

The model was implemented in MATLAB R2025b using the SimBiology toolbox and simulated with the *ode15s* solver, with both absolute and relative tolerances set to 10^*−*12^.

The Sobol’ sensitivity analysis was performed using the *sbiosobol* function and by setting parameter ranges as for the calibration^35^.

For model training, we employed *scattersearch* as the global optimization algorithm (maximum 2000 iterations), followed by *lsqnonlin* as the local solver (maximum 500 iterations). Whenever data variability was available, we performed a weighted fitting with the weights set to the the reciprocal of the variance. The identifiability analysis was conducted with GenSSI^65^, using the mRNA-encoded therapeutic time series as observable.

## Data and code availability

All data supporting the findings described in this manuscript are available in the paper and in the list of references. The Simbiology MATLAB code is available to download at https://github.com/cosbi-research/Campanile_et_al_PBPKmRNATherapeutics.

## Acknowledgments

E.C. is member of the Gruppo Nazionale Calcolo Scientifico, Istituto Nazionale di Alta Matematica (GNCS-INdAM).

## Author contributions

Conceptualization: E.C., E.P., and L.M. Data curation: E.C., E.P., L.L., and L.M. Formal analysis: E.C., S.G., and L.M. Funding acquisition: L.M. Investigation: E.C. and E.P. Methodology: E.C., E.P., S.G., and L.M. Project administration: L.M. Resources: L.M. Software: E.C. and E.P. Supervision: L.M. Validation: E.C., E.P., and L.M. Visualization: E.C., E.P., and L.M. Writing – original draft: E.C., E.P., S.G., L.L., and L.M. Writing – review & editing, E.C., E.P., S.G., L.L., and L.M.

## Declaration of Generative AI and AI-assisted technologies in the writing process

During the preparation of this work the authors used ChatGPT-5 for language refinement and grammar checking. After using this tool, the authors reviewed and edited the content as needed and take full responsibility for the content of the publication.

