## Supplementary material for "Physiologically Based Pharmacokinetic Modeling of mRNA-Encoded Therapeutics: A Multiscale Framework for LNP and Antibody Trafficking in Mice"

Table S1: **95% confidence intervals for the fitted parameters of each product.** Confidence intervals were computed for the best-fit parameter estimates obtained for each product using a Gaussian approximation method.

| | Confidence Intervals | | |
|---|---|---|---|
| | $k_{\text{chain}}$ | $k_{\text{transfer}}$ | $k_{\text{translate}}$ |
| B7H3×CD3 bispecific T-cell engager | [0.10636, 0.38537] | [0.04589, 0.22528] | [59.569, 152.87] |
| RiboMab02.1 | [0.036503, 0.043195] | [0.16552, 0.20974] | [480.46, 588.8] |
| XA-1 | [0.0020032, 0.040488] | [0.0048987, 0.024849] | [41.263, 81.336] |
| Pembrolizumab | [0.0073373, 0.053262] | [0.017534, 0.054468] | [42.625, 62.137] |
| Trastuzumab | [0.031762, 0.15571] | [0.047967, 0.15395] | [19.257, 26.774] |

Table S2: **AAFE values.** Absolute average fold error (AAFE) values computed separately for each therapeutic using model simulations of *(i)* the recombinant therapeutic fit, *(ii)* the mRNA-encoded therapeutic fit, and *(iii)* the mRNA-encoded therapeutic validation.

| | Recombinant therapeutic fit | mRNA-encoded therapeutic fit | mRNA-encoded therapeutic validation |
|---|---|---|---|
| B7H3×CD3 bispecific T-cell engager | 1.6406 | 1.7837 | 1.1064 |
| RiboMab02.1 | 1.8741 | 1.152 | 1.5783 |
| XA-1 | 1.3857 | 1.42 | 1.1466 |
| Pembrolizumab | 1.0947 | 1.374 | 1.241 |
| Trastuzumab | 1.2118 | 1.0583 | 1.2542 |

Table S3: **Parameter value ranges used for calibrations and sensitivity analysis.** Calibration ranges for therapeutic trafficking parameters ($k_{\mathrm{d}}$, $k_{\mathrm{up}}$, and $I_{\mathrm{adj}}$) are derived by considering their minimum and maximum values from the literature. Similarly, ranges for $k_{\mathrm{rec}}$ and $k_{\mathrm{deg}}$ are defined for the sensitivity analysis purpose only. The mRNA-related parameter ranges for $k_{\mathrm{chain}}$, $k_{\mathrm{transfer}}$, and $k_{\mathrm{translate}}$ were set in view of their structural global identifiability and refined through multiple calibration trials to ensure that fitted values remained away from boundary limits.

| Parameter | Unit | Calibration range | Sources and Notes |
|---|---|---|---|
| $k_{\mathrm{d}}$ | Molarity | $[6.30 \times 10^{-8}, 4.77 \times 10^{-5}]$ | Table S2 for details |
| $k_{\mathrm{up}}$ | 1/hour | $[3.22 \times 10^{-2}, 30]$ | Table S3 for details |
| $I_{adj}$ | - | $[0.5, 1]$ | Wiig *et al.* (2008),[1] Sepp *et al.* (2019)[2] |
| $k_{\mathrm{rec}}$ | 1/hour | $[0.28, 11.7]$ | Ferl *et al.* (2005),[3] Sepp *et al.* (2019),[2] Liu *et al.* (2023)[4] |
| $k_{\mathrm{deg}}$ | 1/hour | $[26.1, 152.8]$ | Ferl *et al.* (2005),[3] Sepp *et al.* (2019),[2] Liu *et al.* (2023),[4] Kumar *et al.* (2024)[5] |
| $k_{\mathrm{chain}}$ | 1/hour | $[10^{-3}, 10]$ | - |
| $k_{\mathrm{transfer}}$ | 1/hour | $[10^{-3}, 10]$ | - |
| $k_{\mathrm{translate}}$ | 1/hour | $[1, 10^5]$ | Parhiz *et al.* (2024),[6] Miyazawa *et al.* (2024)[7] |

Table S4: $k_d$ **values from the literature.** Summary of dissociation constant $k_d^{1:1}$ values collected from the literature. When multiple estimates are reported within a single source, the corresponding minimum and maximum values are provided as a range. When only kinetic parameters $k_{on}$ and $k_{off}$ are available, the dissociation constant is calculated following the following formula from Sepp *et al.* (2019):[2] $k_d^{1:1} = k_{off} * avidity / k_{on}$ where $avidity = 10$.

| Source | Value or range of $K_d$ (M) | Notes |
|---|---|---|
| Shah & Betts (2012);[8] Liu & Shah (2023)[4] | $8.12 * 10^{-7}$ | Value computed from $k_{on}$ and $k_{off}$. |
| Liu *et al.* (2024)[9] | $7.5 * 10^{-6}$ | Value computed from $k_{on}$ and $k_{off}$ for endogenous IgG. |
| Jones *et al.* (2020)[10] | $[1.82, 4.77] * 10^{-5}$ | Values obtained for six mAbs at different pH levels. A logarithmic mean was calculated across pH conditions. |
| Abdiche *et al.* (2015);[11] Sepp *et al.* (2019)[2] | $[63, 341] * 10^{-9}$ | Values obtained for IgG1, IgG2, IgG3, and IgG4 subclasses. |

Table S5: $k_{up}$ **values from the literature.** Values of the pynocytosis uptake rate constant $k_{\mathrm{up}}$ from the literature. When multiple estimates are reported within a single source, the corresponding minimum and maximum values are provided as a range.

| Source | Value or range of $k_{\mathrm{up}}$ (1/h) | Notes |
|---|---|---|
| Shah & Betts (2012)[8] | $3.66 * 10^{-2}$ | Reported $\mathrm{CL_{up}}$ values for therapeutic antibodies. |
| Liu & Shah (2023)[4] | $[1, 30]$ | Values including a pinocytosis constant to represent inter-Ab differences. |
| Li *et al.* (2021)[12] | 1.22 | Reported uptake rates in preclinical models. |
| Kumar *et al.* (2024)[5] | 0.32 | Reported uptake rates for mAb PBPK modeling. |
| Sepp *et al.* (2019)[2] | 0.9 | Reference PBPK model parameters for IgG trafficking. |

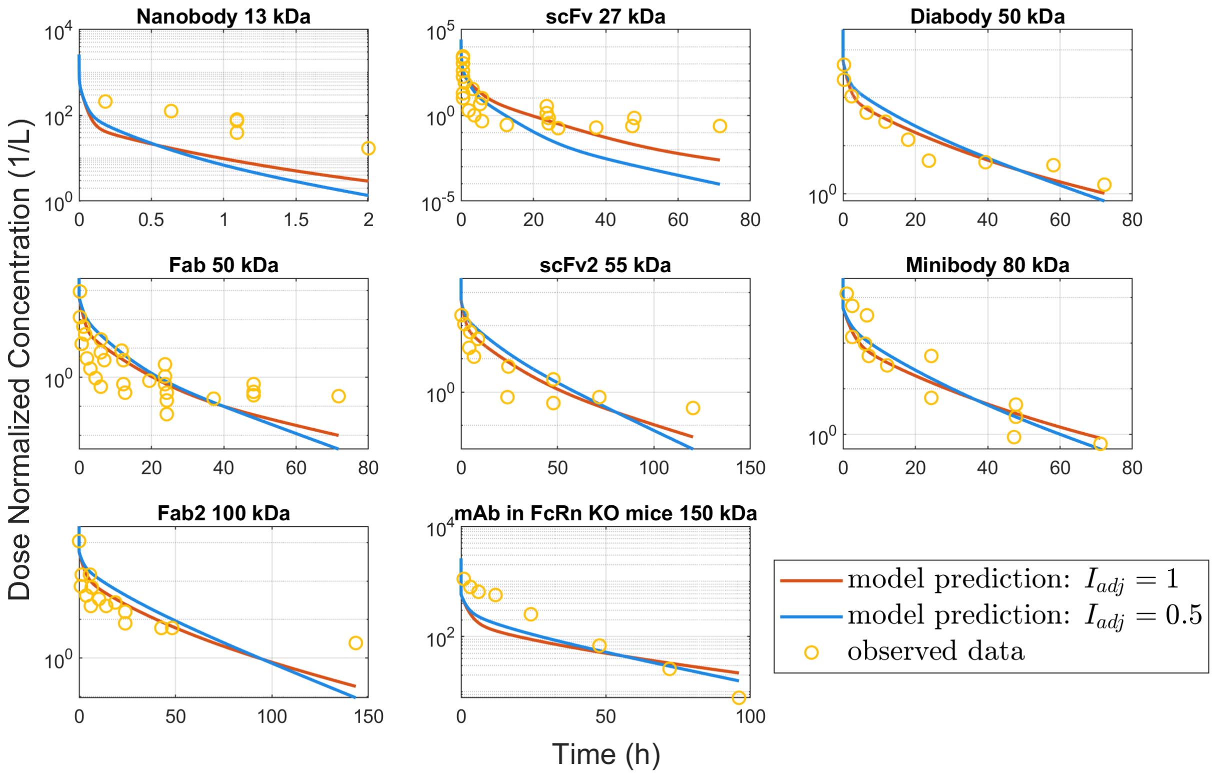


Figure S1: **Calibration plots for eight recombinant therapeutics of different molecular sizes, simulated using either the full or a halved interstitial volume.** For each molecule (13–150 kDa), all lacking FcRn binding, the parameter $k_{\text{up}}$ was fitted to experimental plasma data from Li&Shah (2019)[13] (yellow circles). Model predictions are shown assuming either a full interstitial space ($I_{\text{adj}} = 1$, orange curves) or a halved interstitial space ($I_{\text{adj}} = 0.5$, blue curves). Overall, simulations using the full interstitial volume better reproduce the kinetics of smaller constructs, whereas for larger products—particularly the 80 kDa minibody and the 150 kDa mAb in FcRn-KO mice—the halved-volume assumption yields closer agreement with the observed data.

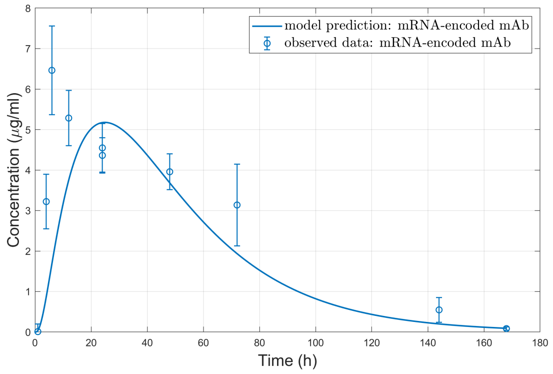


(a) B7H3×CD3 Bispecific T-cell Engager

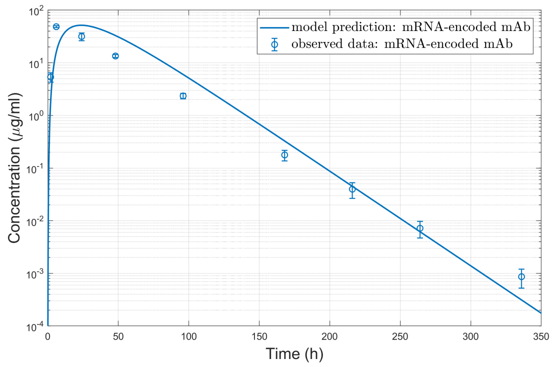


(b) RiboMab02.1

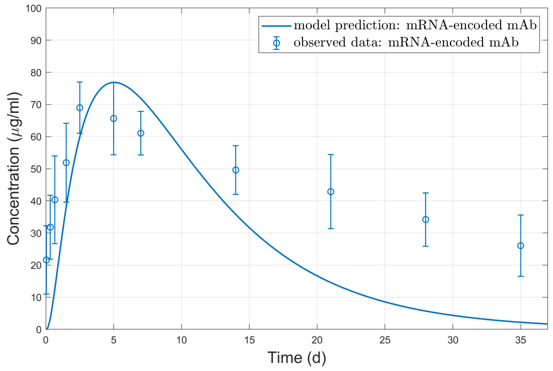


(c) XA-1

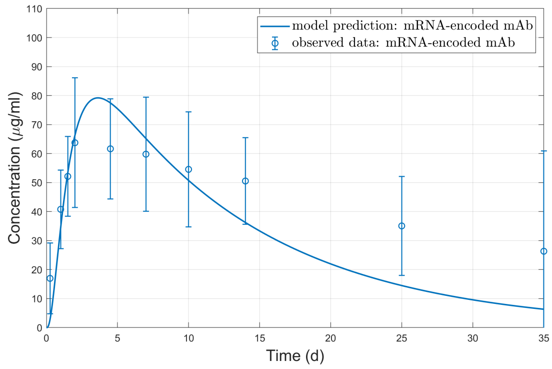


(d) Pembrolizumab

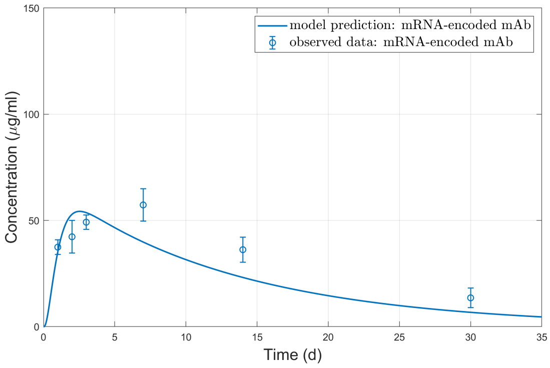


(e) Trastuzumab

Figure S2: **Calibration of the model parameters using a 0-compartment chain.** Calibration time series for each mRNA-encoded therapeutic. The fitting was performed on the mRNA-specific parameters $k_{\text{transfer}}$ and $k_{\text{translate}}$, with $k_{\text{chain}}$ fixed to 0 $h^{-1}$, using the same datasets and dosages as those shown in the left panels (blue curves) of Figure 4 in the main manuscript.

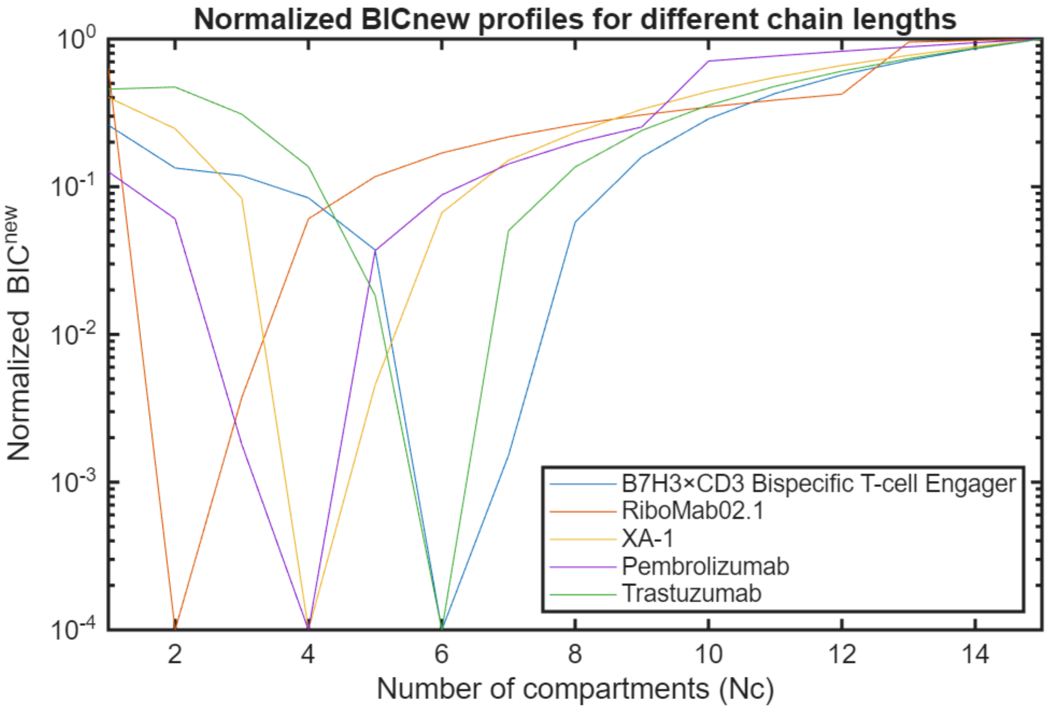


Figure S3: **Normalized $BIC^{\text{new}}$ profiles for different chain lengths across all therapeutics.** For each product, $BIC^{\text{new}}$ was computed for $N^c = 1, \ldots, 15$ and normalized between its minimum and maximum values over the tested range. A small offset of $10^{-4}$ was added for visualization on a logarithmic scale.
